# Severe anaemia complicating HIV in Malawi; multiple co-existing aetiologies are associated with high mortality

**DOI:** 10.1101/666743

**Authors:** Minke HW Huibers, Imelda Bates, Steve McKew, Theresa J Allain, Sarah E. Coupland, Chimota Phiri, Kamija S. Phiri, Michael Boele van Hensbroek, Job C Calis

**Affiliations:** Global child health group, Emma Children’s Hospital, UMC Amsterdam, location Academic Medical Centre, University of Amsterdam, The Netherlands.; Amsterdam Institute of Global Health Development (AIGHD), Amsterdam, the Netherlands.; Liverpool School of Tropical Medicine, Liverpool, United Kingdom.; Department of Internal Medicine, Shrewsbury and Telford Hospital NHS Trust (SaTH), Shrewsbury, United Kingdom; Department of Internal Medicine, College of Medicine, Queen Elizabeth Central Hospital, Blantyre, Malawi.; Department of Molecular and Clinical Cancer Medicine, Institute of Translational Medicine, University of Liverpool, Liverpool, United Kingdom.; Dept. of Pathology, Royal Liverpool University Hospital, Liverpool, UK; School of Public Health and Family Medicine, College of Medicine, Blantyre, Malawi; Department of Pediatric Intensive Care, Emma Children’s Hospital, Academic Medical Centre, University of Amsterdam, The Netherlands.; Department of Paediatrics, College of Medicine, Queen Elizabeth Central Hospital, Blantyre, Malawi.

## Abstract

**Background:** Severe anaemia is a major cause of morbidity and mortality in HIV-infected adults living in resource-limited countries. Comprehensive data on the aetiology is lacking and needed to improve outcomes.

**Methods:** HIV-infected adults with severe (haemoglobin ≤70g/l) or very severe anaemia (haemoglobin ≤50 g/l) were recruited at Queen Elizabeth Central Hospital, Blantyre, Malawi. Fifteen potential causes of severe anaemia of anaemia and associations with anaemia severity and mortality were explored.

**Results:** 199 patients were enrolled: 42.2% had very severe anaemia and 45.7% were on ART. Over two potential causes for anaemia were present in 94% of the patients; including iron deficiency (55.3%), underweight (BMI<20: 49.7%), TB-infection (41.2%) and unsuppressed HIV-infection (viral load >1000 copies/ml) (73.9%). EBV/CMV co-infection (16.5%) was associated with very severe anaemia (OR 2.8 95% CI 1.1-6.9). Overall mortality was high (53%; 100/199) with a median time to death of 16 days. Death was associated with folate deficiency (HR 2.2; 95% CI 1.2-3.8) and end stage renal disease (HR 3.2; 95% CI 1.6-6.2).

**Conclusion:** Mortality among severely anaemic HIV-infected adults is strikingly high. Clinicians must be aware of the urgent need for a multifactorial approach, including starting or optimising HIV treatment; considering TB treatment, nutritional support and attention to potential renal impairment.

## Introduction

Anaemia is recognized as the most common haematological complication of Human Immunodeficiency Virus (HIV) infection worldwide (1, 2). The World Health Organization (WHO) defines anaemia as a haemoglobin level below 110-120 g/l. In sub-Saharan Africa, 60% of HIV-infected adults are anaemic and 22% are severely anaemic (3, 4). Anaemia is independently associated with an increased one-year mortality in HIV infection of 8%, which rises to 55% in those with severe anaemia (5, 6).

To prevent and treat severe anaemia in HIV-infected patients, a comprehensive understanding of the aetiology and pathophysiology is essential. Severe anaemia in HIV infection has been associated with micronutrient deficiencies, infections (viral, bacterial and parasitic), medication induced (zidovudine and co-trimoxazole) and neoplastic diseases (7–11). Only a few studies have comprehensively studied the multifactorial aetiology and pathophysiology of HIV-associated severe anaemia in sub-Saharan Africa despite the high burden of HIV infection in this region (2). Commonly, studies only report on the association between HIV infection and a single cause of severe anaemia, for example iron deficiency, without considering the multiple causes of severe anaemia that may impact on an HIV-infected patient (12, 13). As a consequence, evidence to inform preventive or treatment guidelines for severe anaemia in HIV-infected patients in sub-Saharan Africa is scarce. In practice, severe anaemia management in HIV-infected patients is often still based on the same strategies used for non-HIV infected patients, including iron supplementation, malaria treatment and de-worming (2, 14). This strategy may be ineffective as the causes of HIV-associated severe anaemia may be different, and even harmful considering that (blind) iron supplementation may exacerbate infections and thus may deteriorate the patients condition (15, 16).

Better knowledge about the aetiology of severe anaemia is essential in order to develop evidence-based protocols and ultimately improve outcomes for HIV-infected adults with severe anaemia in sub-Saharan Africa. To address this knowledge gap, we performed a comprehensive observational cohort study to explore the prevalence of potential aetiologies of HIV-associated severe anaemia in Malawian in-patients and studied associations between these and the severity of the anaemia and patient outcomes.

## Methods

An observational cohort study of HIV-infected patients with severe anaemia (haemoglobin ≤70 g/l) admitted to the Queen Elizabeth Central Hospital (QECH), Blantyre, Malawi between February 2010 and March 2011 was performed. All HIV-infected patients above 18 years of age with severe anaemia admitted to the general medical ward were approached and enrolled in the study if they provided informed consent. A case record form including a detailed medical history and physical examination was completed for each enrolled patient. On admission a venous blood sample was collected and a chest X-ray was performed. The patients were managed according to the hospital protocols, which included a blood transfusion if required, treatment of correctable conditions including anti-malarial medication and antibiotics. Antiretroviral treatment (ART) was provided according to the national Malawi guidelines, which stipulated that ARTs should only be initiated by specialist outpatient clinicians, so patients were not initiated on in-patient wards. At the time of the study first line ART was a combination of Stavudine, Lamivudine and Nevirapine and second line treatment was a combination of Zidovudine, Lamivudine, Tenofovir and Lopinavir/Ritonavir (17). ART was only prescribed for WHO stage 3 and 4 disease, or WHO stage 1 and 2 disease with CD4 count < 350 × 10^9^/l (18). Co-trimoxazole was prescribed routinely to all HIV-infected patients on ART as Pneumocystis Jirovecii prophylaxis. For this study patients were followed up in the dedicated ART clinic after discharge. A community research nurse followed those who failed to attend their appointments up at home. Follow-up was done for a maximum of 365 days after enrolment when they attended the ART clinic for routine appointments or, if they failed to attend, by a home visit from a study nurse.

### Laboratory assays

All samples were analysed within 24 hours of collection or stored at −80°C for further analysis. On enrolment haemoglobin concentrations were measured on the ward using the HemoCue B-Haemoglobin analyser (HemoCue, Ängelholm, Sweden) to screen for eligibility. For patients enrolled in the study, the haemoglobin and red cell indices (MCV, MCH were determined and MCHC) using an automated analyser (Beckman Coulter, Durban, South Africa). CD4-cell counts were assessed using BD FACS Count (BD Biosciences, San Jose, CA, USA). Transferrin, iron, ferritin, folate and vitamin B12 were analysed on Modular P800 and Monular Analytic E170 systems (Roche Diagnostics, Switzerland). Soluble transferrin receptor (sTfR) levels were measured using ELISA (Ramco Laboratories, TX, USA). Analyses for serum creatinine was by Beckman Coulter CX5 (ADVIA 2400 Siemens Healthcare Diagnostics). Renal function was measured by a estimating of Glomerular Filtration Rate (eGFR) using simplified Modification of Diet in Renal Disease (MDRD)-Study formula and the GFR was classified by Chronic Kidney Disease classification (19–21). For all tests the manufacturers’ reference ranges were used; internationally accepted cut-offs were used to define deficiencies (22). Thick blood films were prepared and stained for malaria microscopy. Malaria was defined as the presence of *Plasmodium falciparum* asexual parasites in the blood films. HIV infection was confirmed using two point-of-care antibody tests (Unigold® and Determine®). A venous blood sample was inoculated into BACTEC Myco/F-Lytic culture vials and incubated in a BACTEC 9050 automated culture system (Becton Dickinson) for 56 days. Sub-culturing blood and sputum, susceptibility testing and isolate identification were performed by standard techniques(23). Possible contaminants were recorded as absence of pathogens. Sputum cultures were examined for mycobacteria using Ziehl–Nielsen staining. Whole-blood isolates were assessed for Epstein–Barr virus and cytomegalovirus infection by semi-quantitative PCR and for parvovirus B19 by real-time PCR (24). All chest X-rays were reviewed by a radiologist for signs of pulmonary tuberculosis (TB). `When TB was suspected, standardized treatment was started according to the local protocols.

### Bone marrow

If the patient’s clinical condition allowed and they provided consent, a bone marrow aspirate and trephine biopsy was performed. All bone marrow samples were taken from the posterior iliac crest. Samples of the aspirates were spread onto slides and trephine biopsies were fixed, decalcified and embedded in paraffin wax (25, 26). Bone marrow samples were sent to the Haematopathology Referral Centre at the Royal Liverpool University Hospital, Liverpool UK, for analysis. Sections of the trephine blocks were stained with haematoxylin and eosin and Giemsa, and also for iron with Perls stain, and for reticulin (25). All slides were examined for using a predefined format and diagnoses were allocated to the categories lymphoproliferative disease, myeloproliferative disease, myelodysplastic syndrome (MDS) and TB (27–29). The need for additional histochemical (e.g. Ziehl-Neelsen) or immunohistological (e.g. CD3, CD20) staining was determined according to the local protocol in Liverpool depending on the preliminary morphological findings.

### Definitions for potential factors involved in the aetiology of severe anaemia

The following potential factors involved in the aetiology of severe anaemia were defined and evaluated; 1) Unsuppressed HIV-infection; viral load ≥1000 copies/ml. 2)TB: one or more of the following were present: a) positive sputum culture, b) chest X-ray with signs of pulmonary TB and/or c) on going TB treatment at time of enrolment d) clinical diagnosis by local doctor including unknown generalized lymphadenopathy and/or night sweats of > 30 days and of unknown origin e) caseating granulomata in the bone marrow trephine. 3) Malaria: presence of malaria parasites in a thick blood film. 4) Parvovirus B19: viral load of >1000 copies/ml. 5) Cytomegalovirus (CMV); load of >100 copies/ml. 6) Epstein-Barr virus (EBV); viral load >100 copies/ml. 7) Bacteraemia; a blood culture growing a potential pathogen. 8) Underweight (BMI ≤18.5). 9) Serum folate deficiency (≤3 ng/l). 10) Vitamin B12 deficiency (≤180 pg/ml). 11). Iron deficiency was defined as MCV≤ 83 fl (3, 22, 30). 12) Zidovudine usage. 13) Co-trimoxazole usage. 14) Bone marrow disorders; lympho-proliferative disease, myeloid-proliferative disease or MDS. 15) Renal impairment: a GFR which either indicated impaired (GFR 15–59 ml/min/1.73 m^2^) or End Stage (GFR ≤15 ml/min/1.73 m^2^) Renal Disease (21, 31).

### Statistics

Baseline characteristics and prevalence of potential risk factors are presented as proportions or medians with IQR. Logistic regression was performed to model the association between anaemia and potential factors involved severity of anaemia, Results are expressed as OR with 95% CI and p-values. Variables associated with the outcome variables (*P* ≤ 0.10) in the univariate analysis were included in the multivariate model in a stepwise approach. Kaplan Meier survival curves were used to assess cumulative mortality. Significant differences were investigated with a Log Rank test. Uni- and multivariate analyses were done using logistic regression and Cox regression to describe predictors of mortality. P values of ≤ 0.05 were regarded as statistically significant. All reported P values were two-sided. The data were analysed using Stata (version 12) (STATA Corp. LP, Texas, TX, USA).

### Ethics

The Research Ethics Committee of the College of Medicine, University of Malawi (P.09.09.824) and the Research Ethics Committee of Liverpool School of Tropical Medicine (research protocol 09.64) approved the study. The purpose of the study was explained to the patients in the local language Chichewa and written informed consent was obtained before inclusion in the study.

## Results

In total, 199 patients were included in the study: 64.8% were female. The median age was 32 years (IQR 27-61 years). The median haemoglobin was 53 g/l (IQR 4.2-6.3) and 84 (42%) patients had very severe anaemia (hb≤50 g/l). A total of 91 (45.7%) patients were on ART at enrolment including 79.1% on first line ART. During the study period, an additional 41 (21%) patients started on ART treatment. Moreover 67.1% of the patients were immune suppressed with a CD4 count ≤ 200 cells/mm. Baseline characteristics of the patients are shown in table1.

**Table 1.** Baseline characteristics of HIV-infected patients with severe anaemia at enrolment into study. ^*1*^Pancytopenia is defined as thrombocytopenia (≤150×10^9^/l) and leukopenia (≤4×10^9^/l) and severe anaemia (≤ 70 g/l) (22). Abbreviations: Hb: Haemoglobin, ART: antiretroviral therapy.

|  | Overall |
| --- | --- |
| Age, years (median IQR) | 32 (IQR 27-61) |
| Sex (female) | 129/199 (64.8%) |
| **Haematology** |  |
| Haemoglobin (Hb) (median, IQR) g/l | 53 (IQR 42-63) |
| Severe anaemia (Hb 51-70 g/l) | 115/199 (57.8%) |
| Very severe anaemia ($\text{Hb} \leq 50 \text{ g/l}$) | 84/199 (42.2%) |
| Pancytopenia<sup>1</sup> | 42 /184 (21.2%) |
| **Mortality** |  |
| Overall mortality (365 days) | 101/199 (50.8%) |
| Early mortality (60 days) | 81/101 (81.2%) |
| Days until death (median, IQR) | 17.5(6-55) |
| **HIV-disease and treatment** |  |
| CD4 (median, IQR) | 175 (IQR 55-825) |
| $\text{CD4} \leq 200 \text{ cells/mm}^3$ | 104/155 (67.1%) |
| Viral load $\geq 1000 \text{ copies/ml}$ | 136/184 (73.9%) |
| ART at enrolment | 91/199 (45.7%) |
| First-line ART | 72/199 (36.2%) |
| Second-line ART | 11/199 (5.5%) |
| Non-specified ART | 8/199 (4%) |
| **Blood transfusions and supplemental treatment** |  |
| Blood transfusion at admission | 47/199 (23.6%) |
| Folate supplementation at admission | 57/199 (28.6%) |
| Iron supplementation at admission | 81/199 (40.7%) |
| Vitamin B12 supplementation at admission | 0/199 (-) |

The prevalence of factors that were associated with severe anaemia, and factors that were co-existing in individual patients, are shown in table 2. Overall, patients had a mean of 3 (range 1-8) co-existing aetiologies of severe anaemia (figure 1).

**Table 2.** Distribution and multivariate analysis of co-existing factors associated with severe and very severe anaemia (Hb≤ 50 g/l) in HIV-infected adults in Malawi. ^*1*^A total of 28-blood cultures were positive, the most common organisms were E. coli (42.9%; 12/28) and non-Typhoid Salmonella (17.9 %; 5/28). Explanatory variables associated with the outcome variables (P > 0.10) in the univariable analysis were excluded in the multivariable model in a stepwise approach (-). Abbreviation: GFR; Glomerular filtration rate.

|  | Overall (n=199) | Severe anaemia<br>N=115/199<br>(30.8%) | Very severe<br>anaemia<br>N=84/199<br>(42.2%) | Univariate |  |  | Multivariate |  |  |
| --- | --- | --- | --- | --- | --- | --- | --- | --- | --- |
|  |  |  |  | Odds | 95% -CI | P-value | Odds | 95% -CI | P-value |
| Sex (female) | 129 (64.8%) | 74/115 (65.2%) | 55/84 (65.4%) | 1.1 | 0.6-1.9 | 0.869 | - | - | - |
| **HIV** |  |  |  |  |  |  |  |  |  |
| CD4 ≤200 cells/mm<sup>3</sup> | 104/155 (67.1%) | 62/87 (71.3%) | 42/68 (61.8%) | 1.04 | 0.6-1.8 | 0.889 | - | - | - |
| Viral load ≥1000 copies/ml | 136/184 (73.9%) | 81/104 (77.9%) | 55/80 (68.8%) | 0.6 | 0.3-1.2 | 0.163 | 0.7 | 0.3-1.5 | 0.306 |
| On ART at enrolment | 91/199 (42.7%) | 52/115 (40.9%) | 39/84 (45.2%) | 1.1 | 0.6-1.8 | 0.865 | - | - | - |
| **Infection** |  |  |  |  |  |  |  |  |  |
| Malaria | 6/167 (1%) | 3/100 (3.0%) | 3/67 (4.5%) | 1.5 | 0.3-7.8 | 0.617 | - | - | - |
| Tuberculosis | 82/199 (41.2%) | 53/115 (46.0%) | 29/84 (34.5%) | 0.7 | 0.6-0.99 | 0.043 | 0.6 | 0.1-2.8 | 0.507 |
| Bacteraemia<sup>1</sup> | 26/199 (13.1%) | 17/115 (61.7%) | 9/82 (11.0%) | 0.7 | 0.3-1.6 | 0.402 | - | - | - |
| Parvovirus B19 | 7/170 (4.2%) | 5/99 (5.0%) | 2/71 (2.8%) | 0.5 | 0.1-2.9 | 0.476 | - | - | - |
| Cytomegalovirus (CMV) | 57/170 (33.5%) | 28/99 (28.3%) | 29/71 (40.8%) | 1.8 | 0.9-3.3 | 0.088 | - | - | - |
| Epstein-Barr virus (EBV) | 69/170 (40.6%) | 40/99 (40.4%) | 29/71 (40.8%) | 1.0 | 0.6-1.9 | 0.954 | - | - | - |
| EBV/CMV co-infection | 28/170 (16.5%) | 12/99 (12.1%) | 16/71 (22.5%) | 2.1 | 0.9-4.5 | 0.075 | 2.8 | 1.1-6.9 | 0.032 |
| **Malnutrition** |  |  |  |  |  |  |  |  |  |
| Underweight | 74/148 (49.7%) | 40/81 (49.3%) | 34/68 (50.0%) | 1.0 | 0.5-2.0 | 0.940 | - | - | - |
| Vitamin B12 deficiency | 2/194 (1.0%) | 1/113 (0.8%) | 1/81 (1.2%) | 1.4 | 0.09-22.7 | 0.813 | - | - | - |
| Iron deficiency | 61/180 (33.9%) | 38/104 (36.5%) | 23/76 (30.3%) | 0.8 | 0.40-1.42 | 0.380 | - | - | - |
| Folate deficiency | 23/194 (11.9%) | 13/113 (11.5%) | 10/84 (11.9%) | 1.1 | 0.5-2.6 | 0.858 | - | - | - |
| **Medication** |  |  |  |  |  |  |  |  |  |
| Co-trimoxazole | 163/199 (81.9%) | 99/115 (81.6%) | 64/84 (76.2%) | 0.5 | 0.3-2.3 | 0.076 | 0.5 | 0.2-1.2 | 0.099 |
| Zidovudine | 13/199 (6.5%) | 5/50 (10.0%) | 8/40 (20.0%) | 2.3 | 0.7-7.5 | 0.187 | - | - | - |
| **Renal function** |  |  |  |  |  |  |  |  |  |
| Normal (GFR >60) | 149/185 (80.5%) | 90/105 (85.7%) | 59/80 (73.8%) | 1 |  |  | 1 |  |  |
| Impaired (GFR 15-60) | 24/185 (13.0%) | 12/105 (11.4%) | 12/80 (15.0%) | 1.5 | 0.6-3.6 | 0.339 | 1.0 | 0.7-5.0 | 0.176 |
| End stage (GFR ≤15) | 12/185 (6.5%) | 3/105 (2.9%) | 9/80 (11.3%) | 4.6 | 1.2-17.6 | 0.027 | 3.0 | 0.7-13.1 | 0.140 |
| **Bone marrow** |  |  |  |  |  |  |  |  |  |
| Bone marrow disease | 28/71 (39.4%) | 19/42 (45.2%) | 9/29 (31.0%) | 0.5 | 0.2-1.5 | 0.231 | - | - |  |
| **Aetiology** |  |  |  |  |  |  |  |  |  |
| Co-existing aetiologies per patient (mean, SD) | 3.3 (1.3) | 3.3 (1.2) | 3.2 (1.4) | 0.8 | 0.4-1.8 | 0.605 | - | - | - |

**Figure 1.**
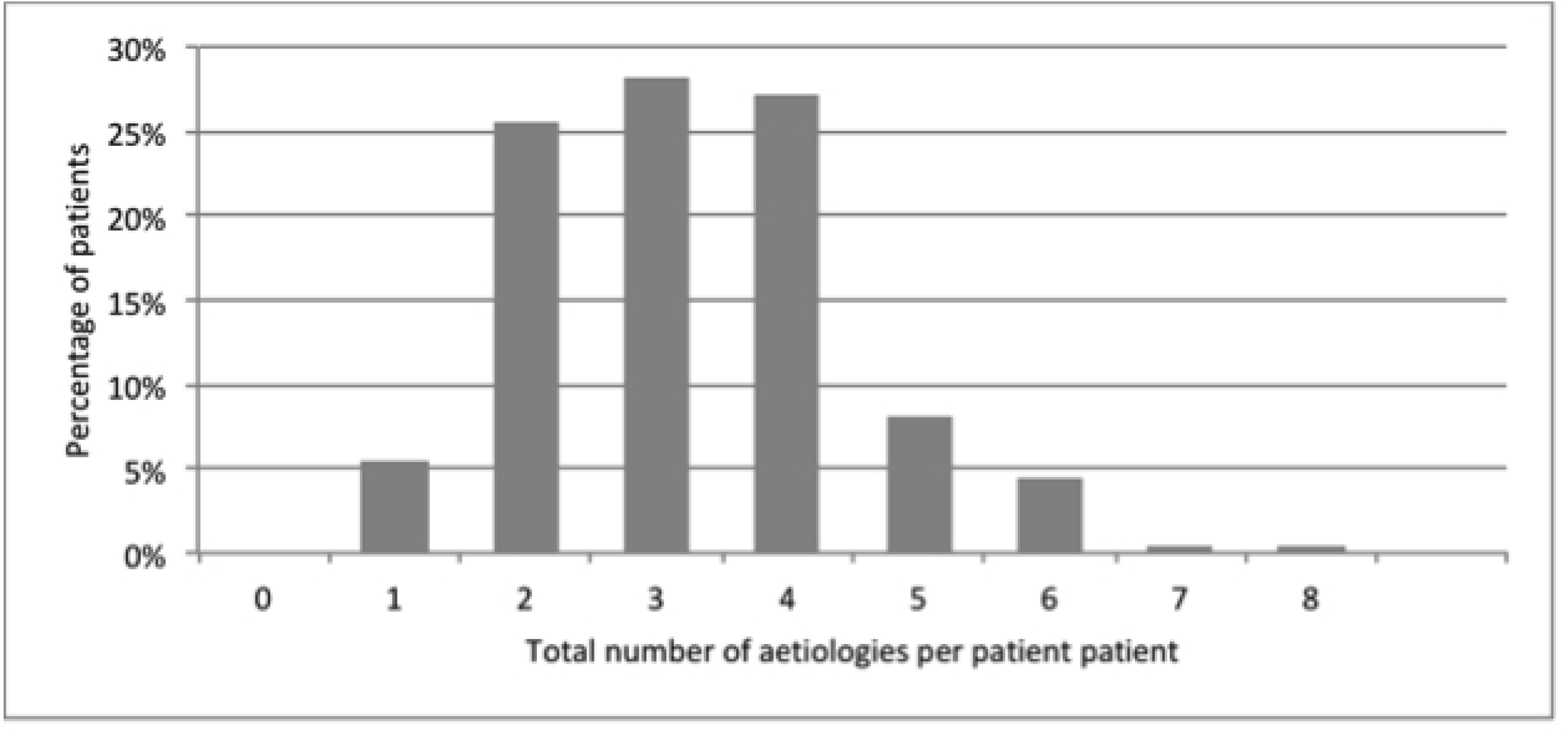
Total number aetiologies for severe anaemia co-existing in each patient (n=199). Mean is 3 factors (SD 1.3), range 1-8.

An unsuppressed HIV-infection was seen among 73.9% patients. TB was the second most common infection occurring in 82 (41.2%) of the patients. In 19 (23%) of these patients TB was diagnosed on their chest X-ray. Granulomata were seen in the bone marrow trephine in 15 (18%) patients of all the 82 patients that had a diagnosis of TB. 11 (13%) patients were on TB treatment at enrolment. 69/170 (40.5%) patients had evidence of current EBV infection and 57/170 (33.5%) had evidence of current CMV infection. Co-infection with CMV and EBV was found in of 28/170 (16.5%) of the patients. Bacteraemia was diagnosed on a positive blood culture in 26 (13.1%); the most common pathogens found were E. Coli (12 patients; 42.9% and non-Typhoid Salmonella (5 patients; 19.9%). 74/148 (49.7%) patients were underweight and iron deficiency occurred in 61/180 (33.9%) of the patients. Bone marrow sampling was performed in 73 patients. Of these, 28 (38.4%) had morphological abnormalities with MDS being the most common abnormality (20 patients; 27.4%) (table 2). Renal impairment was diagnosed in 36/185 patients (19.5%) and 12 of these patients (33%) had end stage renal disease.

Comparing the different risk factors for very severe anaemia (Hb<50g/L) as compared to severe anaemia, EBV/CMV co-infection (OR 2.8 95% CI 1.1-6.9) was the only factor associated with very severe anaemia (table 2).

During the one-year follow-up period, 101 study patients (50.8%) died. The median time to death was 16 days (IQR 5-41) and 81 (80.1%) of these deaths occurred within 60 days of admission (figure 2).

**Figure 2.**
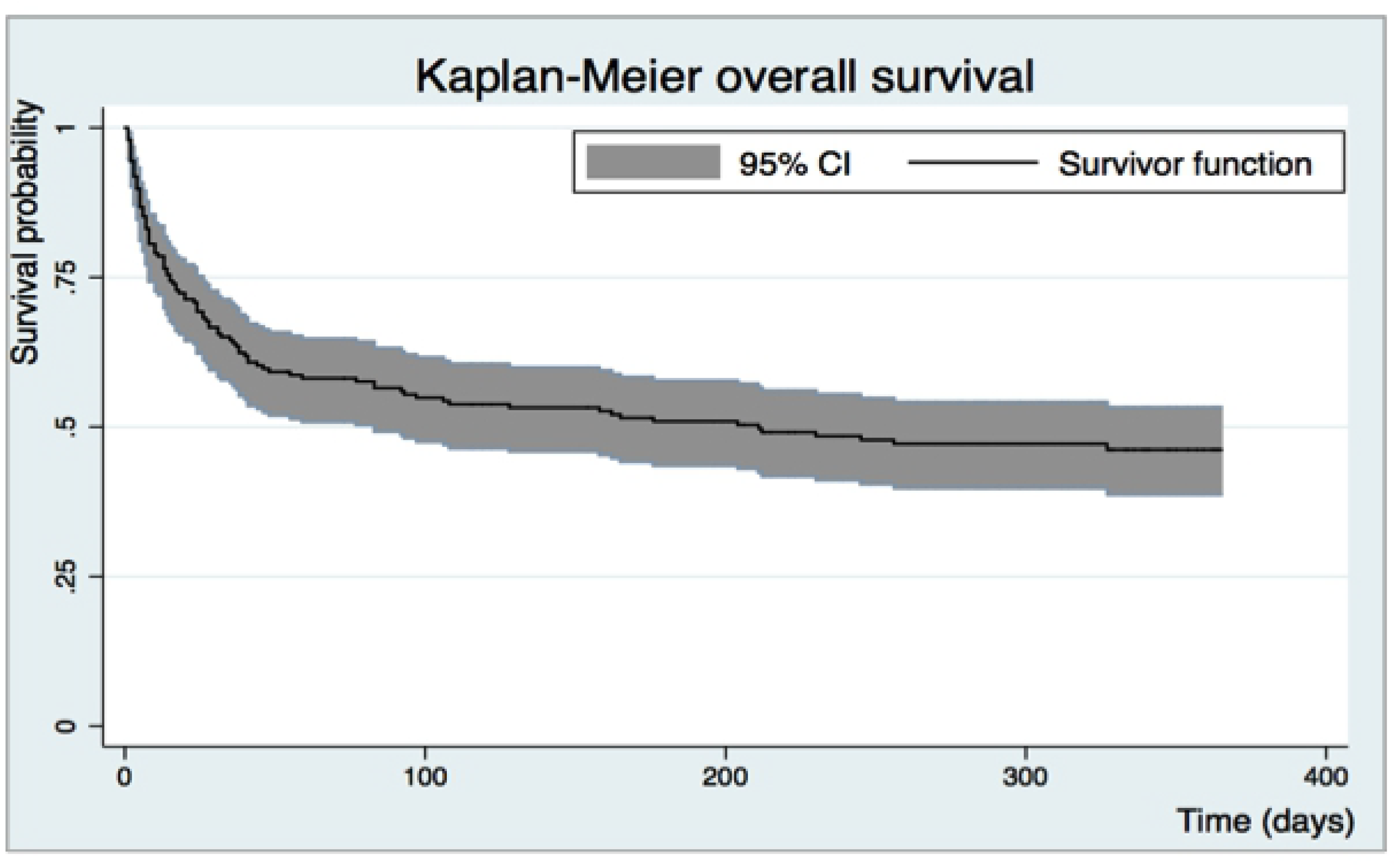
Kaplan Meyer curve for survival probability over time (days) for adult Malawian patients with HIV infection and severe anaemia during 365 days follow-up. Abbreviation: 95% confidence interval (95%CI).

Folate deficiency and end stage renal disease were associated with mortality with Hazard Ratio 2.2 (95% CI 1.2-3.8) and Hazard Ratio 3.2 (95% CI 1.6-6.2) respectively (figure 3). Neither very severe anaemia (haemoglobin ≤50 g/l), nor the haemoglobin levels were associated with mortality in the study patients (Hazard Ratio 0.9, 95%CI 0.6-1.4 and Hazard Ratio 1.01, 95% CI 0.9-1.2 respectively).

**Figure 3.**
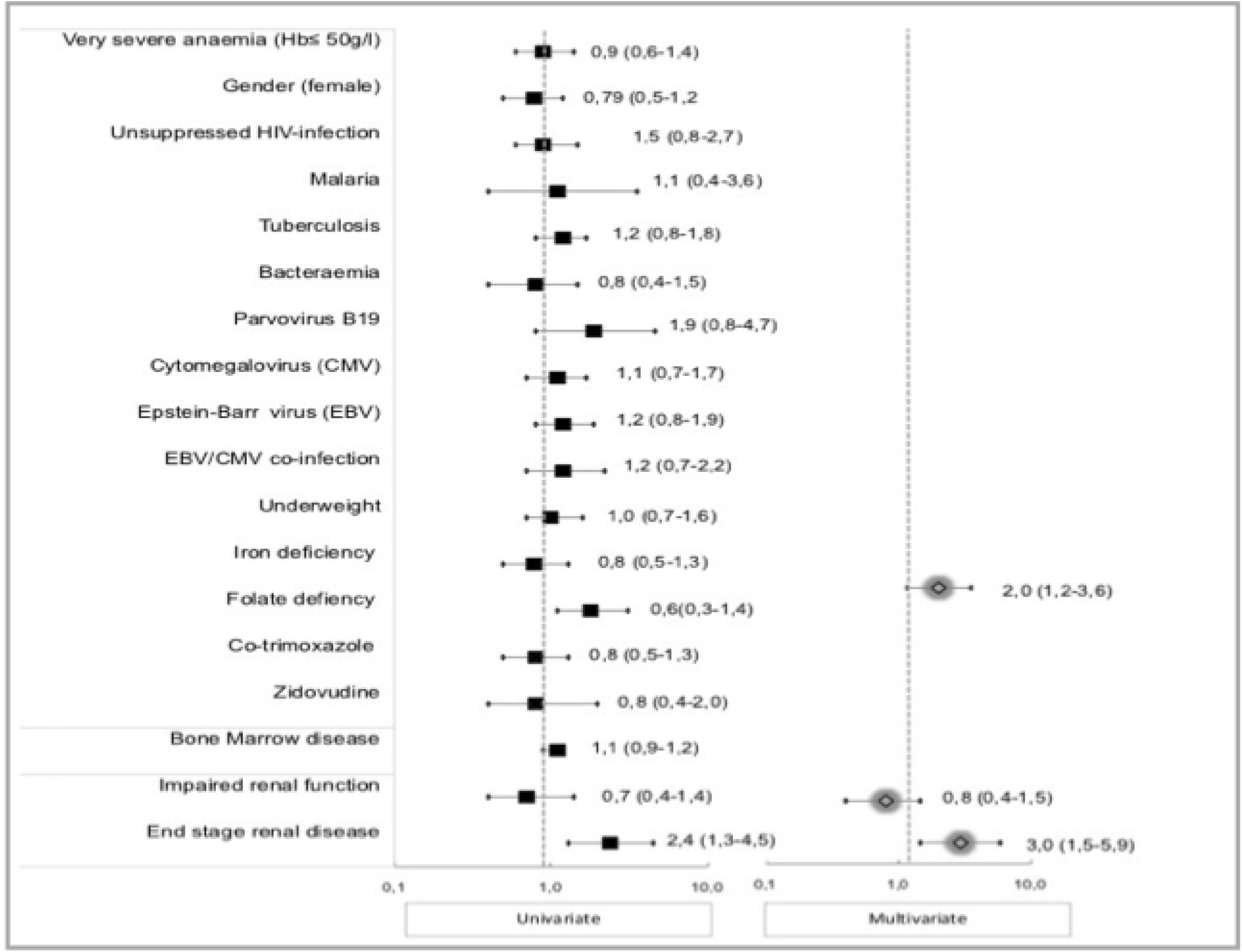
Risk factors for 365-day mortality in HIV-infected patients with severe anaemia. Univariate and multivariate Cox regression outcome. Folate deficiency (3 ng/l); HR 2.1 95% CI 1.2-3.6 and end stage renal disease (GFR ≤15); HR 3.2 5% CI 1.6-6.2, were associated with overall mortality. Abbreviations: Hb: Haemoglobin, GFR; Glomerular filtration rate, VL Viral Load.

## Discussion

In this study we described the prevalence of several potential aetiologies for severe and very severe anaemia in HIV-infected Malawian adults. Patients had a mean of three co-existing aetiologies potentially contributing to their anaemia, the most common one being unsuppressed HIV. Mortality in the study patients was extremely high, as 51% of the patients died within one year and most died within 60 days of admission. Severe anaemia in HIV-infected patients in a resource limited setting, such as Malawi, should therefore be treated as a multi-causal critical condition with high mortality.

Anaemia is an independent predictor of mortality and disease progression in HIV-infected patients and mortality increases with decreasing haemoglobin concentration (6, 32). Previous studies reported an estimated one-year mortality of 3.7% in HIV-infected patients without anaemia and up to 30-55% in severe anaemia in resource limited settings (5, 6). Our study results are consistent with these outcomes. Additionally we found that a mean of three contemporaneous aetiologies potentially contributed to severe anaemia in HIV-infected patients. This is important as it highlights the multi-causality of severe anaemia in this population. Clinicians should therefore not just focus on a single cause and single treatment for severe anaemia in HIV-infected patients. Different treatment protocols from those used traditionally to treat anaemia (e.g. iron and folate) will be needed to address these multiple co-existing aetiologies to enhance improvement of mortality amongst these patients. Also, the seriousness of severe anaemia in HIV-infected patients needs to be better recognized. Currently severe anaemia in HIV-infected patients is classified as a stage 3 clinical condition (33). However due to the high mortality found in our, and previous, studies, we recommend re-classifying severe anaemia as a marker of HIV stage 4 disease.

Unsuppressed HIV virus was present in 79% of the patients in our study population. HIV may cause anaemia directly through an inhibitory effect of the HIV-virus on the erythropoietin progenitor cells in the bone marrow, or indirectly through opportunistic infections causing anaemia (34). Although this potential direct/indirect risk factor was very common it was not associated with the severity of the anaemia or mortality in our population, explored in our multivariate analysis. This is in contrast with a large cross-sectional study in Tanzania showed that the risk of developing severe anaemia in HIV-infected patients increased two to three times among patients with advanced HIV disease (3). We therefor postulate that controlling HIV infection by starting or switching ART treatment should therefore be considered as the most important and urgent step in treatment protocols for severely anaemic HIV-infected patients. After our study had been completed guidelines for initiating ART changed. At the time of the study the trigger for starting ART was based on a patient’s CD4 count, whereas it is currently recommended that ART should be started early in the course of HIV disease (17, 33). It will be important for the impact of this policy change on HIV-related anaemia, and its consequences, to be evaluated. Irrespective of the policy change for initiating ART, the findings of our study remain very relevant because many HIV-infected patients in resource-limited settings present late in the course of their disease or are unable to access reliable supplies of ART, and are therefore likely to continue to have high levels of life-threatening anaemia.

TB has previously been associated with anaemia in HIV-infected patients (36). In our study patients TB was a common co-infection (41%). This prevalence is comparable to previous reports on TB prevalence (43%) among severely anaemic HIV-infected patients in Africa (8, 14). The pathophysiology of TB-associated anaemia in HIV-infected patients remains unclear. Bone marrow invasion by TB organisms or altered iron metabolism, as a side effect of tuberculostatic drugs have been described (36). Only 13% of patients were on TB treatment at enrolment. TB medication itself was not associated with the severity of anaemia (data not shown). Given that we and others have found TB in nearly half of the HIV-infected patients with severe anaemia, TB screening and rapid initiation of treatment should therefore be a high priority in the management of these patients, especially in resource limiting settings where there is a high TB prevalence.

Viral infections such as EBV, CMV and parvovirus B19 have been associated with anaemia in HIV-infected patients (35, 37). In our study EBV and CMV were common and present in 40% and 35% of patients respectively, whilst parvovirus B19 was less common (4%). Although parvovirus B19 is pathophysiological linked to mild anaemia, the role in the development severe anaemia in HIV infected patients has not been clearly established (38). Data on co-infections are very limited in African patients (39–41) and our study is the first to describe the association between co-existing CMV and EBV infections and very severe anaemia (haemoglobin ≤50 g/l). Possible explanations for this association include direct viral inhibition of erythropoiesis (39, 42, 43). The majority of our patients with co-infection had advanced HIV disease (88.5%) but the association between EBV and CMV co-infection and very severe anaemia remained significant even after correction for advanced HIV disease. As CMV is a treatable infection it will be important to determine the effect of ART on CMV infection and severe anaemia, as ART can reduce CMV infection by improving immune status (44).

Malaria is unsurprisingly associated with severe anaemia in HIV-infected patients in sub-Saharan Africa (2, 14). However, the contribution of malaria to anaemia in our study population was small however, as only 6 patients (3.5%) had malaria parasites on presentation. This is in line with data from other studies that show that the role of malaria in causing anaemia, especially in HIV patients in sub-Saharan Africa (2, 14), is limited and likely to have been overestimated (40, 45).

Malnutrition was common in our population as half of the patients’ had a BMI below 18.5m^2^ and deficiencies of iron and folate were diagnosed in 33.9% and 12% of the patients respectively. The prevalence of iron deficiency in our study is higher than in previous studies on HIV-infected patients in sub-Saharan Africa populations that report prevalence rates of iron deficiency of 18%-25% of anaemic HIV-infected patients (46, 47). Interestingly, adults in sub-Saharan Africa with severe anaemia but without HIV infection, have a higher prevalence of iron deficiency (59%) (45). Data from children with severe anaemia in Malawi also showed a much higher overall prevalence of iron deficiency (47%) than the adults in our study (40). Studies presenting this outcome are all published before 2010, in this time period ART was less available and all included HIV-infected patients were ART naïve and sever immune suppression was highly present. In this group of HIV-infected patients with better immune systems the aetiology of severely anaemic may be more similar to the aetiology of non-HIV infected patients. It can be expected that the prevalence of condition such as iron deficiency is also more alike the HIV-uninfected groups (40). In addition to finding a high prevalence of iron deficiency, ours is the first study to document on a possible association between folate deficiency and increased mortality in severely anaemic HIV-infected patients. There are a limited number of studies describing micronutrient supplementation, including folate, for anaemic HIV-infected adults, but overall there appears to be little significant effect of supplementation on reducing morbidity and mortality (37, 48). Macronutrient support using, for example, fortified wheat flower, has had beneficial effects on anaemia reduction and micronutrient levels in populations in sub-Saharan Africa, but has never been tested in the context of severe anaemia in HIV-infected patients (49, 50). More research is therefore needed to evaluate the effectiveness of both macro- and micro-nutritional support in severely anaemic HIV-infected adults.

HIV-infected adults have an increased risk of neoplastic bone marrow diseases, which can often cause anaemia (13). However, prevalence data for such conditions among HIV-infected patients with anaemia, especially in resource limiting settings, are scarce. Only three of our study patients had confirmed bone marrow malignancies. In contrast MDS was common occurring in 27% of our patients (27, 51).

Renal impairment was a frequent finding among our study patients and has been linked to HIV disease progression, anaemia and poor outcomes in both wealthy and resource limited settings (52–55). Evaluation of renal function is an important component of severe anaemia treatment protocols (30, 53) since it may affect the choice of ART (52, 56). It may also have implications for clinical management, for example by introducing measures to prevent further deterioration or considering potential benefits of erythropoietin(53).

Our study has several limitations. We purposely did not explore factors potentially associated with anaemia that previous studies had shown were uncommon in Malawi, such as haemoglobinopathies and parasitic infections (40, 57). We also did not include an HIV-infected population without severe anaemia against which to compare the prevalence of anaemia aetiologies and clinical outcomes. Only a sub-population of our patients gave consent for bone marrow sampling. Although these were an unselected group of patients it is possible that this may have introduced a hidden bias, for example by excluding the patients who were particularly unwell. Nevertheless, our findings are very valuable since bone marrow data from HIV-infected African patients is very scarce.

## Conclusion

Our study has demonstrated that severe anaemia in HIV-infected adults in Malawi is associated with multiple co-existing aetiologies and has a strikingly high mortality rate. Severe anaemia in HIV-infected patients is therefore a critical indicator of mortality and requires urgent and multiple interventions. Particularly important is the initiation of ART, the management of infections such as TB and CMV, and optimisation of renal function. Intervention studies are needed to properly define the role (and safety) of iron and folate supplementation, as well as to develop and evaluate guidelines, which are feasible and effective in resource-limited settings to help clinicians manage these patients more effectively.

## Acknowledgement

The authors would like to thank all of the study participants, doctors, nurses and support staff of Queens Elizabeth Hospital and the Malawi-Liverpool-Wellcome centre in Blantyre for their participation and cooperation. This study was supported by the Nutricia research foundation (Project number 2017-43), The Hague, the Netherlands and the Wellcome Trust (Project number WT086559), Liverpool, United Kingdom. The funders had no role in the study design, data collection and analysis, decision to publish or preparation of the manuscript

